# Haplotype differences in common bean accessions confer the capacity to flower under Scandinavian summer conditions

**DOI:** 10.1101/2023.08.17.553676

**Authors:** M Rendón-Anaya, G Buinovskaja, L Yu, PK Ingvarsson

**Affiliations:** Linnean Centre for Plant Biology, Department of Plant Biology, Uppsala BioCenter, Swedish University of Agricultural Sciences, Uppsala, Sweden

## Abstract

The capacity to flower under long days has been a major pre-requisite for the adaptation of the common bean to European climates. The complexity of such adaptation has been studied, mostly under the optics of QTL mapping, but still the genetic basis of the trait remains elusive. In the current study we sequenced a collection of >200 accessions of *P. vulgaris* of Mesoamerican, Andean and European origin, for which the flowering capacity under long days during the summer season in Sweden was evaluated. Our variant calling strategy allowed us to identify 16.9e^6^ SNPs and 38.6e^3^ long structural variants. Furthermore, we observed gene-pool specific selective sweeps that correspond to the independent domestication events in the Americas. GWAS and haplotype structure tests identified single nucleotide and structural variants strongly associated to the capacity to flower under long days, particularly in chromosome 1.

## Introduction

Many aspects of plant growth and development show fine-tuned local adaptation, and this is particularly true for the transition from vegetative growth to flowering (Roux, et al. 2006). In many plant species, flowering requires exposure to specific photoperiods and/or temperatures and flowering may be delayed or prevented when these requirements are not met. As plants encounter environments that subject them to novel conditions, genetic adaptations that modify, relax or eliminate existing constraints on flowering can enable expansion across environmental gradients and/or climatic regimes. Although flowering time can be seen from one perspective as a relatively simple trait, the transition from vegetative to reproductive stages is accompanied by significant changes to a wide range of other developmental traits, including stem elongation, apical dominance, lateral branching, and resource allocation. Thus, a large number of genes that contribute to the control of flowering time have been identified in *Arabidopsis* and other model plant species (Huang and Nusinow 2016; Huang, et al. 2017), and it has been observed that genetic networks regulating flowering and the transitions between vegetative and reproductive stages are largely conserved across Angiosperms (Pin and Nilsson 2012).

Crops domesticated in the Americas span a spectrum of genetic relatedness, have been domesticated for diverse purposes, and have responded to human selection by changes in many different traits. While these crops provide examples of both parallel and convergent evolution at various levels, data are still insufficient to provide quantitative or conclusive assessments of the relative roles of these two processes in domestication and diversification. The common bean (*Phaseolus vulgaris* L) as a species originated in Mesoamerica, which is also the centre of origin for the entire genus *Phaseolus. P. vulgaris* is a herbaceous annual plant that mainly reproduces through selfing. Two *P. vulgaris* gene-pools have been recognized, one in Mesoamerica and the second in the southern Andean region in South America, and these gene pools diverged around 165,000 YA (Schmutz, et al. 2014; Ariani, et al. 2018). Common bean was then independently domesticated in these two locations around 8,000 YA (Kwak and Gepts 2009), leading to two distinct populations that share similar morphological traits as a consequence of artificial selection. Following the arrival of Spain and Portugal to the Americas, the common bean was introduced to Europe where it rapidly adapted to grow at both higher latitudes and lower temperatures over the last 500 years. The European cultivars of *P. vulgaris* have a predominant Andean genetic background (Angioi, et al. 2010).

Populations of *P. vulgaris* comprise either short-day plants that flower under short-day photoperiods (day length less than 12hrs) that allow seed production to be completed before the onset of the dry season, or day-neutral plants that flower independently of photoperiod cues. Following the introduction to Europe, common bean has been under selection for altered photoperiod sensitivity in favour of flowering under long day conditions (day lengths >14 hrs) which coincides with the growing season across much of Europe (latitudes 35°N and higher). Today, European landraces show clear signs of adaptation to local photoperiod (Rodriguez, et al. 2013). However, such a rapid response to the European local conditions could also be mediated by pre-existing adaptations to cooler climates and differences in photoperiod sensitivity in the introduced genotypes.

Genetic analyses of flowering in legumes, e.g., soybean, pea and alfalfa, have associated dozens of relevant genes to photoperiod sensitivity, including genes involved in light perception, the circadian clock or signal integration for inflorescence development [reviewed by (Weller and Ortega 2015)]. In common bean, recombinant inbred lines (RIL) based analyses have associated several loci to photoperiod response, e.g., PHYTOCHROME A (Weller, et al. 2019) on chromosome 1, a locus in chromosome 4 encoding a CONSTANS-like and a gene encoding an AGAMOUS-like 8 on chromosome 9 (Gonzalez, et al. 2021).

In this report we sequenced the full genome of a large collection of 232 *P. vulgaris* accessions from different geographic origins, including wild individuals, landraces and elite cultivars. We dissected the signatures of domestication and adaptation through scans of positive selection and studied the genomic background of the flowering capacity during long days by means of GWAS and haplotype structure analyses. The resolution of our comparisons allowed us to identify genomic signals specific to each gene-pool and to identify different haplotype structures that allow common bean cultivars to flower under long days, typical of the Scandinavian summer season.

## Materials and methods

### Plant material and growing conditions

We collected a diverse panel of 232 *Phaseolus vulgaris* accessions of Mesoamerican, Andean and European origin, that include commercial accessions, land races and wild-collected individuals from the centres of origin (Supplementary table 1). These accessions are publicly available at the International Centre for Tropical Agriculture in Cali, Colombia; the Nordic Genetic Resource Center and the European Search Catalogue for Plant Genetic Resources at Gatersleben, Germany.

We evaluated our panel of *Phaseolus* accessions as follows: the 232 accessions were sown in medium pots (10 cm of diameter) with 750 gr of sterile soil. Plants were grown in green-house conditions during the summer at the Plant Cultivation Facility at the BioCentre, Swedish University of Agricultural Sciences, Uppsala, which means that plants were exposed to very long days (>16hrs of light) and temperatures as high as >28°C. The capacity to flower under these conditions was coded as a binary trait (0, non-flowering; 1, flowering) for the association analyses.

### Genome re-sequencing, mapping and SNP calling

Total genomic DNA was extracted from frozen leaf tissue for all individuals using the DNeasy plant mini prep kit (QIAGEN, Valencia, CA, USA). Briefly,1 μg of high-quality DNA was used for paired-end libraries construction. The libraries (TruSeq, PCR-free, 350bp) were subjected to paired-end sequencing (2×150) at the National Genomics Infrastructure at Science for Life Laboratory, Stockholm, on an Illumina NovaSeq 6000 to a mean per-sample depth of approximately 15X.

Raw reads were mapped to the reference genome of *P. vulgaris* (https://phytozome-next.jgi.doe.gov/info/Pvulgaris_v2_1) using BWA-mem version 2.2.3 (Li and Durbin 2009). Aligned reads were flagged for duplicates using the MarkDuplicates program in Picard tools version 1.119 (http://broadinstitute.github.io/picard/). Multi-sample single nucleotide polymorphism (SNP) calling was performed using the Genome Analysis Toolkit (GATK) version 3.8. For GATK, SNPs were called using the tools HaplotypeCaller and GenotypeGVCF. SNPs were only retained if they matched the following criteria: QualByDepth > 20 (a measure of alternative allele quality independent of read depth), MQ > 30 (the minimum phred-scaled mapping quality of the reads supporting the alternative allele), FisherStrand < 10 (whether an alternative allele was predominantly supported by one read orientation only),−2 < BaseQualityRankSumTest < 2 (a test statistic to assess whether the base quality of reads supporting the alternative allele was significantly worse than reads supporting the reference allele), −2 < ReadPosRankSumTest < 2 (a test statistic to assess whether the base position in a read supporting the alternative allele was significantly different than the base position in a read supporting the reference allele),−2 < MappingQualityRankSumTest < 2 (a test statistic to assess whether the mapping quality of reads supporting the alternative allele was significantly worse than reads supporting the reference allele), and RMSMappingQuality > 30 (an estimation of the overall mapping quality of reads supporting an alternative allele).

We calculated genome-wide breadth and depth of coverage using samtools (Li, et al. 2009). After obtaining the average coverage in our collection, we set a minimum and maximum number of reads per site to filter out very low/high depth sites. For downstream analyses we kept only biallelic sites with no more that 30% missingness in the vcf file, which were obtained with BCFtools (Li 2011).

### Population genomics

To calculate standard population genomics estimators, such as pi, F_ST_ and dxy we used the python package *genomics_general* (https://github.com/simonhmartin/genomics_general). Linkage-disequilibrium (LD) decay was plotted with PopLDdecay (Zhang, et al. 2019). In order to remove potentially duplicated samples between genebanks and closely related individuals we calculated identity by descent (IBD) with PLINK v1.9 (Purcell, et al. 2007). Samples with a PI_HAT > 0.4 were removed. SNP-based PCAs were constructed using plink –pca on pruned, unlinked sites with a minimum MAF exceeding 0.05.

We used two methods to assign the ancestral allelic state to each polymorphic site. We used est-sfs (Keightley and Jackson 2018), that estimates the unfolded site frequency spectrum, allowing for several outgroups and nucleotide substitutions models. To use this method, we selected two *P. coccineus* genotypes as outgroups, one of them domesticated (G35406) and a wild, Mesoamerican (G35856); we ran estsfs using the rate-6 model. However, mapping *P. coccineus* on the reference *P. vulgaris* genome results in a reduced breadth of coverage (∼70% of *P. coccineus* vs 92% of *P. vulgaris*). For this reason, we also produced a consensus genome based on the most represented variants in the wild Mesoamerican *P. vulgaris* samples using ANGSD -doFasta 2 that outputs a fasta file taking the most common base per site given a series of bam infiles mapped to the reference genome (Korneliussen, et al. 2014). With this, we could extrapolate the ancestral allele in the vcf files for further selective sweep analyses.

### Structural variant identification

We combined three different software to detect structural variants in the samples, DELLY (Rausch, et al. 2012), dysgu (Cleal and Baird 2022) and Manta (Chen, et al. 2015). Our in-house pipeline finds SVs supported (flagged PASS) by the three software with minimum 80% reciprocal overlap, and produces a single VCF file with the filtered, high-quality SVs (https://github.com/buinovsg/SV_detection_pipeline/). Because of the amount of memory required by each software prevented analyses of large numbers of samples, we called SVs in four batches, MW, AW, MD+AD and EU, and then combined them in one single VCF file assuming genotypes at missing types as reference with bcftools merge -0 (Li 2011). We searched for gene-pool specific SVs assuming a minor allele frequency greater than 0.05 to avoid spurious signals.

### Positive selection scans

Scans for selective sweeps were performed using both a cross-population composite likelihood ratio method (XP-CLR, implemented in python (Chen, et al. 2010)), and the cross-population extended haplotype homozygosity test (XP-EHH, implemented in the rehh package version 2.0.2 in R (Gautier and Vitalis 2012)) on subsamples identified with our PCA analysis and using only SNPs that were polymorphic within the genetic cluster. The pre-defined subpopulations correspond to Mesoamerican wild (MW), Mesoamerican domesticated (MD), Andean wild (AW), Andean domesticated (AD), European (EU) that can be subdivided in European with Mesoamerican background (Eu-M) or European with an Andean background (Eu-A). These populations will be referred to using their respective abbreviated names. We used pairwise comparisons as follows: wild vs domesticated (MW-MD; AW-AD) and domesticated vs introduced (MD-EuM; AD-EuA), in order to differentiate the effects of the domestication process from recent adaptation to the European environment. XP-EHH assesses haplotype differences between two populations and is designed to detect alleles that have increased in frequency to the point of fixation or near fixation in one of the two populations being compared. XP-CLR, on the other hand, is based on the linked allele frequency difference between two populations and is a unidirectional method to find the pattern with regional allelic frequency difference in-between population. We combined the top 5% of Fst and top 5% XP-CLR as outliers in each pairwise comparisons for a more stringent identification of windows with signatures of positive selection. For XP-EHH, significant SNPs with p-value < 0.05 were considered significant. We used default options for all analyses, setting the window size to 10kb with a minimum of 100 polymorphic sites per window, given that the SNP density per KB is close to 10 in all populations but the MW. Because of the slow decay of LD in the domesticated clusters we next binned chromosomes in 500kb windows and calculated the occurrence of XP-CLR-Fst outliers in each bin. Based on the distribution of windows, we extended windows if the neighbouring bin had at least 5 outlier windows.

### Genome wide associations and haplotype differentiation

We coded the phenotypic data as binary matrices, i.e. flowering accessions were coded with 1, whereas non-flowering were coded with a 0. We used SNP panels of pruned sites (filtered with plink -maf 0.05 -indep-pairwise 100 10 0.2 -geno 0.1) to run genome-wide association analyses. We produced three different sets of pruned SNPs, one for the entire collection of *P. vulgaris* accessions, and two more for the gene-pool specific association analyses. We converted each vcf file to a matrix of 0,1 and 2 values for homozygous (ref/alt) or heterozygous genotypes with vcftools (vcftools –012). As we allowed 10% of missingness in this genotype filtering step, we had to impute the missing genotype information to avoid numeric biases in the GWAS calculations. For each column in the matrix representing individual positions, we calculated the mean genotype value (not including missing genotypes coded as -1) that we used to fill the missing genotypes. This numeric matrix was the input for GWAS analyses.

We combined genotype and phenotypic data on photoperiod sensitivity through genome-wide association mapping (GWAS). GWAS was performed using several models: mixed linear model (MLM), multiple loci mixed model (MLMM), and Bayesian-information and linkage-disequilibrium iteratively nested keyway (BLINK), all implemented in GAPIT3 (Wang and Zhang 2021). We controlled for the effects of population structure by setting the number of relevant PCs at 5; we used SNPs and SVs with a minimum allele frequency of 0.05.

Finally, we conducted haplotype structure analyses around significant markers associated to our target phenotype in the GWAS screenings, which represent a very efficient method to identify differences in LD decay. For these small scale analysis we polarized the variant sites using the est-sfs method. We performed haplotype furcation analyses, where the root of each diagram is a selected marker – defined as a SNP with significant p-values in our GWAS screenings. We also calculated the relative EHH, i.e. the factor by which EHH decays at the focal SNP compared with the decay of EHH on all other core haplotypes combined. EHH at a distance x from a core SNP is defined as the probability that two random chromosomes carrying a tested core haplotype, are homozygous at all variant sites for the entire interval from the core region to the distance x. Then, the haplotype structure was evaluated in a bi-directional mode, allowing us to estimate both proximal and distal LD. Moving in one direction, the diagram might bifurcate if both or only one allele is present at the next marker. The thickness of the lines corresponds to the number of samples with the long-distance haplotype. EHH and bifurcation analyses were run with rehh (R; (Sabeti, et al. 2002; Gautier and Vitalis 2012)). This level or resolution allowed us to see the structure of derived and ancestral haplotypes in the tested populations in contrast to control individuals, such as the wild samples from each species and gene pool.

## Results

### Population structure

We re-sequenced 232 *P. vulgaris* accessions of Mesoamerican, Andean and European origin, that include commercial accessions, land races and wild-collected individuals from the centres of origin (Table S1). The estimated average depth and breadth of coverage were 13,7X and 90.2% across the samples. Due to strong relatedness and to possible duplications of some accessions across seedbanks, we retained 172 accessions for downstream analyses.

Based on 161,391 pruned SNPs out of a total of 16,907,953, we obtained a PCA that grouped the accessions following their genetic background: PC1 separates the accessions in the two dominant gene pools, Mesoamerican (MA) and Andean (AN), while PC2 separates them according to their domesticated/wild state (Figure 1A). We observe two divergent groups of wild accessions, two clusters of domesticated accessions in proximity to their wild ancestors, which confirms both independent domestication events, and finally, a large group of accessions grown in Europe that can be separated according to their genomic background, but that also displays clear signs of admixture following the introduction of the species in Europe. Based on this result, we further separate the European collection according to their predominant genomic background (Table S2).

**Figure 1.**
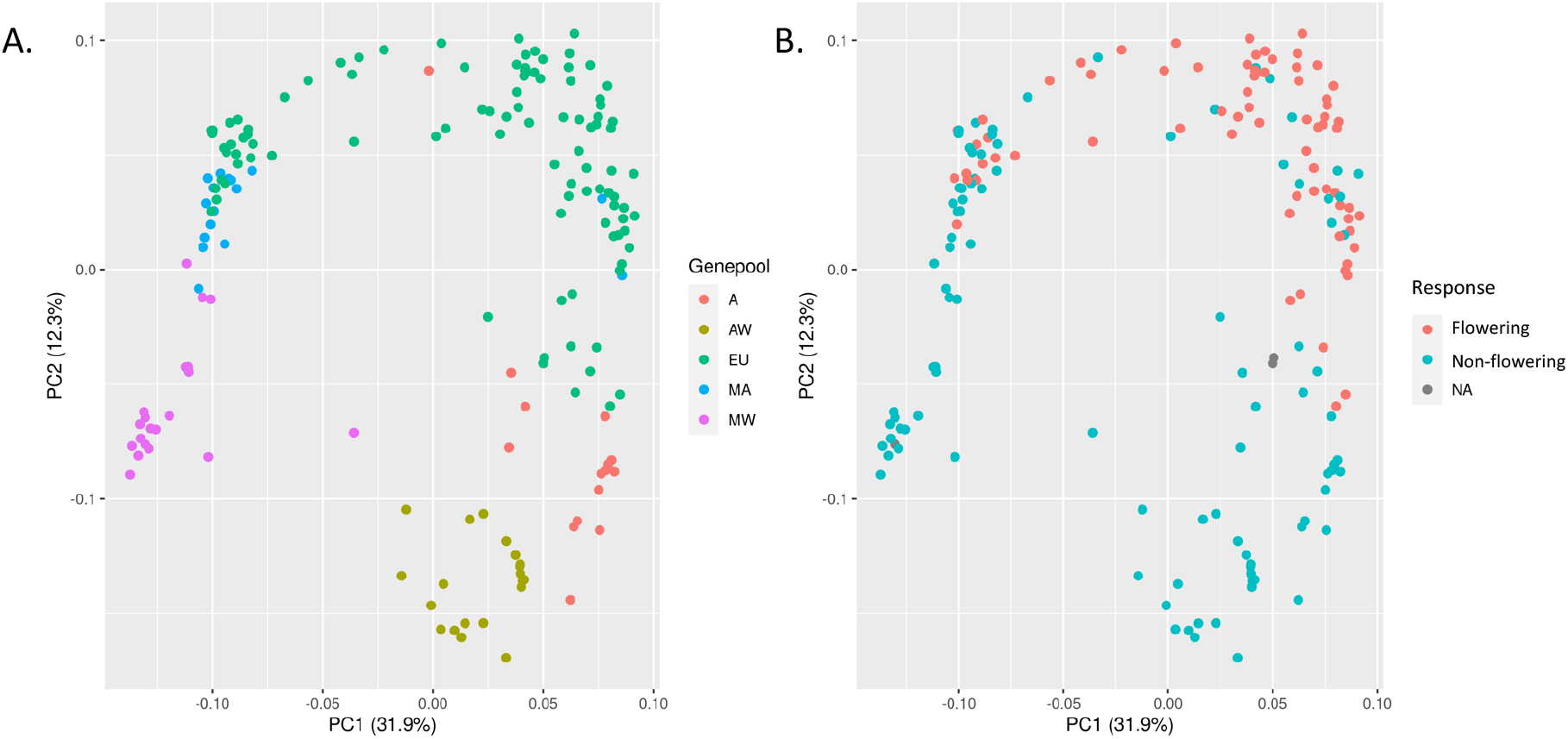
Population structure. SNP-based PCAs were constructed using non-rare, unlinked variants across the 11 chromosomes. The accessions are labeld according to their genetic background (A) or their sensitivity to photoperiod (B).

Interestingly, when classified by flowering capacity (Figure 1B), we confirm the day length sensitivity in the wild accessions. Furthermore, most accessions grown in Europe had a predominant Andean background, which confirms previous observations regarding the gene pools present in the continent (Gepts and Bliss 1988; Angioi, et al. 2010).

### Population genomics

We ran several estimators of genetic diversity and differentiation in our panel of accessions that confirmed previous observations in terms of diversity and linkage disequilibrium decay between gene pools. First, clustering of the accessions based on their dxy values produced two independent clades, one Mesoamerican and one Andean, with longer branch lengths in the MA clade (Figure S1). We observed the highest nucleotide diversity in the wild Mesoamerican population (pi[MW]=0.13; Figure S2A), which is expected given that MA is the centre of origin of the species (Alfonso Delgado-Salinas 2006; Delgado-Salinas, et al. 2006). The lowest diversity was observed in the Andean gene pool (pi[AW]=0.02; pi[AD]=0.025) while in European samples we observed a recovery of diversity which is a likely indicator of the recent admixture between MA and A gene pools following their introduction in Europe. The calculated inbreeding coefficient (F) in each subpopulation followed the expected gradient (from ∼0.7 in MW up to ∼0.96 in AD) given the preferred self-pollination strategy of *P. vulgaris* and its demographic history (Figure S2B). In terms of linkage disequilibrium, we observed a rapid LD decay in less than 50kb in wild MA samples while LD extends several hundreds of Kbs in the domesticated populations and particularly individuals with an Andean background (Figure S3A).

### Positive selection and sweep detection

A selective sweep occurs when a beneficial variant arises and spreads in a population and results in reduced sequence diversity both at the site of the beneficial allele as well as at neutral markers linked to the selected site (Tiffin and Ross-Ibarra 2014). Since selective sweeps result in reduced variation, a distorted site frequency spectrum, high linkage disequilibrium and extended haplotype structure in genomic regions surrounding the sites of fixed adaptive mutations, they are relatively easy to detect using a number of statistical methods (Sabeti, et al. 2002; Ferrer-Admetlla, et al. 2014; Vatsiou, et al. 2016). We used an array of selection statistics to identify regions targeted by positive selection during domestication in the Americas and adaptation to European conditions over the past five centuries. We used both site frequency spectrum (XP-CLR) and linkage disequilibrium (XP-EHH) based approaches on subsamples of Andean and Mesoamerican origin in a pairwise fashion: wild vs domesticated (MW-MD; AW-AD) and domesticated vs introduced (MD-EuM; AD-EuA).

Using a stringent significance threshold (top 5% XP-CLR values on 10Kb windows with at least 100 variants per window given the SNP density per Kb; Figure S3B) we obtained 891 and 224 outlier windows for the MD and AD gene pools, respectively. To achieve an even more stringent filter of selective sweep regions we combined the top 5% XP-CLR signals with the top 5% F_ST_ outliers from each pairwise comparison. The intersection of XP-CLR and F_ST_ outliers resulted in 248 and 100 windows as potential domestication targeted regions in the MA and AD gene pools, respectively (Figure 2A, highlighted in green). Interestingly, we found only four overlapping regions between these MA and AD domestication related regions, in chromosomes 8, 9 and 11 containing the gene models Phvul.008G246000 (homologous to CRF4 cytokinin response factor 4), Phvul.009G057700 (homologous to the DUF21 domain-containing protein At1g47330), Phvul.011G065400 (homologous to a ubiquitin-conjugating enzyme E2 7) and Phvul.011G065500 (homologous to ATDSI-1VOC, a dessication-induced 1VOC superfamily protein). The same filters were applied to the clusters of European accessions (separated by genetic background), obtaining 58 and 13 outlier windows in the EU-MD and EU-AD, respectively (Figure 2B). Because of the slow decay of LD in domesticated samples, we grouped significant windows identified as XP-CLR outliers in bins of 500kB across all chromosomes. In the case of the MA domestication event, we observed a high number of XP-CLR outliers (>=10 windows/500Kb; average number of outliers/500Kb=0.9) in chromosomes 1 between 47.5-48Mb, 12 outliers; chromosome 3, 50.5-51Mb; chromosome 4, 10-10.5Mb; chromosome 7, 2-2.5Mb; chromosome 8, 4.5-5Mb, 58.5-60Mb (25 outliers in total in these 1.5Mb), and chromosome 9, 10.5-11 Mb, 13-13.5Mb and 30.5-31Mb. Such signal was weaker for the AD domestication event (average number of outliers/500Kb=0.2) presumably because of lower power to detect selective sweeps due to the substantially reduced diversity seen across the genomes of plants with an Andean origin.

**Figure 2.**
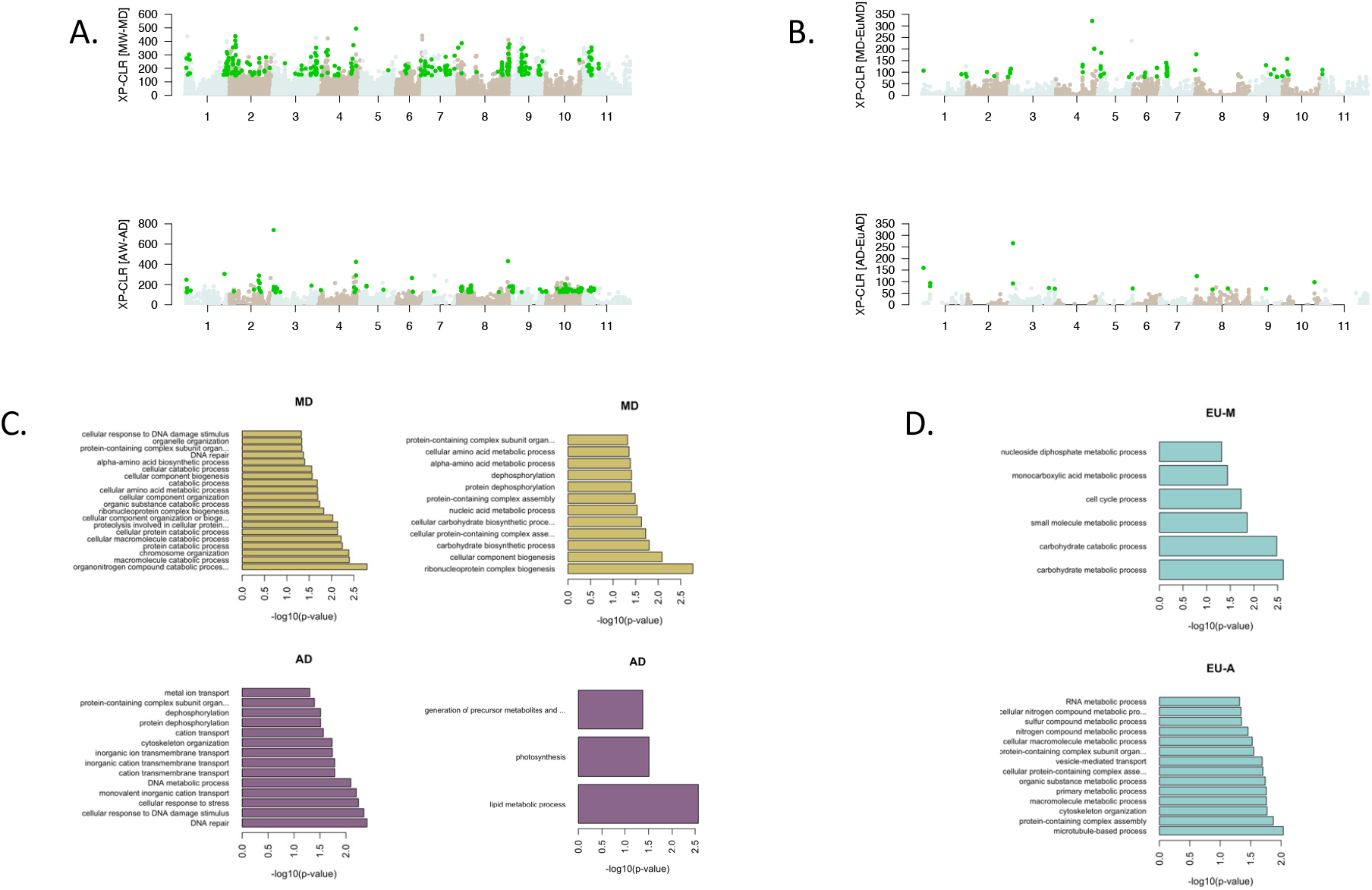
Scans of positive selection during domestication and adaptation. A. XP-CLR screening of positive selection associated to domestication events in Mesoamerica (MW-MD) and the Andes (AW-AD). B. XP-CLR screening of selection associated to the adaptation to Europe. The overlaps of XP-CLR and Fst outliers are highlighted in green. C and D. Functional enrichments in Fst (left) and XP-CLR (right) outlier windows.

Using an LD-based approach of cross-population extended haplotype homozygosity we obtained a different set of signals of domestication and adaptation. This pairwise comparison had a low performance when contrasting domesticated accessions from the Americas against their wild relatives, detecting only 10 and 34 significant sites in AD and MA respectively, at a p-value <0.05 for |XP-EHH|. However, this method detected 277 significant sites when contrasting EUMD against the MA domesticated subsample. Of these, only 2 sites on chromosome 2 overlapped with an XP-CLR outlier in the MA gene pool, but there was no intersection in the AD gene pool cross-comparisons.

We then used the gene models predicted within the outlier windows of our selection scans and evaluated functional categories that were over-represented among them using F_ST_ and XP-CLR outliers separately as the stringent intersection of both statistics did not produce any significant enrichment categories. We observed interesting GO terms significantly enriched in the MA domestication event such as those related to carbohydrate and amino acid metabolic processes. In the AD pool, terms related to lipid metabolic process, photosynthesis/light harvesting, and generation of precursor metabolites and energy were detected. Interestingly, in both domesticated gene pools, DNA repair and cellular response to DNA damage stimulus, as well as protein dephosphorylation were significantly enriched (Figure 2C). In the outliers associated with adaptation to Europe, we observed similar categories such as carbohydrate, nitrogen and sulfur compound metabolic processes (Figure 2D). Only accessions with an MD background were enriched for protein phosphorylation at XP-CLR outliers.

### Structural variation in the domesticated gene-pools

Structural variation (SV) is defined as large genomic differences (> 50 bp) between individuals which arise from changes in DNA sequence length, copy number, orientation, or chromosomal location. They can be classified into deletions (DELs), duplications (DUPs), insertions (INSs), inversions (INVs) and translocations (TRAs).

Most SVs are formed from single events but different combinations of SV types have been known to occur together (Yi and Ju 2018). SVs are usually considered separately from single nucleotide polymorphisms (SNPs) and small indels (insertions and deletions) due to their larger size, greater impact on gene function and different mutational origin (Chiang, et al. 2017). Based on the intersection of three software designed to detect SVs from short read data, we identified 38,591 variants in our panel of accessions. A PCA analysis based on SVs (filtered by MAF>0.05) recovered the population structure we detected previously with SNP data (Figure 3A). As expected, given the Andean origin of the reference genome (Schmutz, et al. 2014), we detected the highest number of unique SVs in the MW pool followed by AW and MD (Figure 3B).

**Figure 3.**
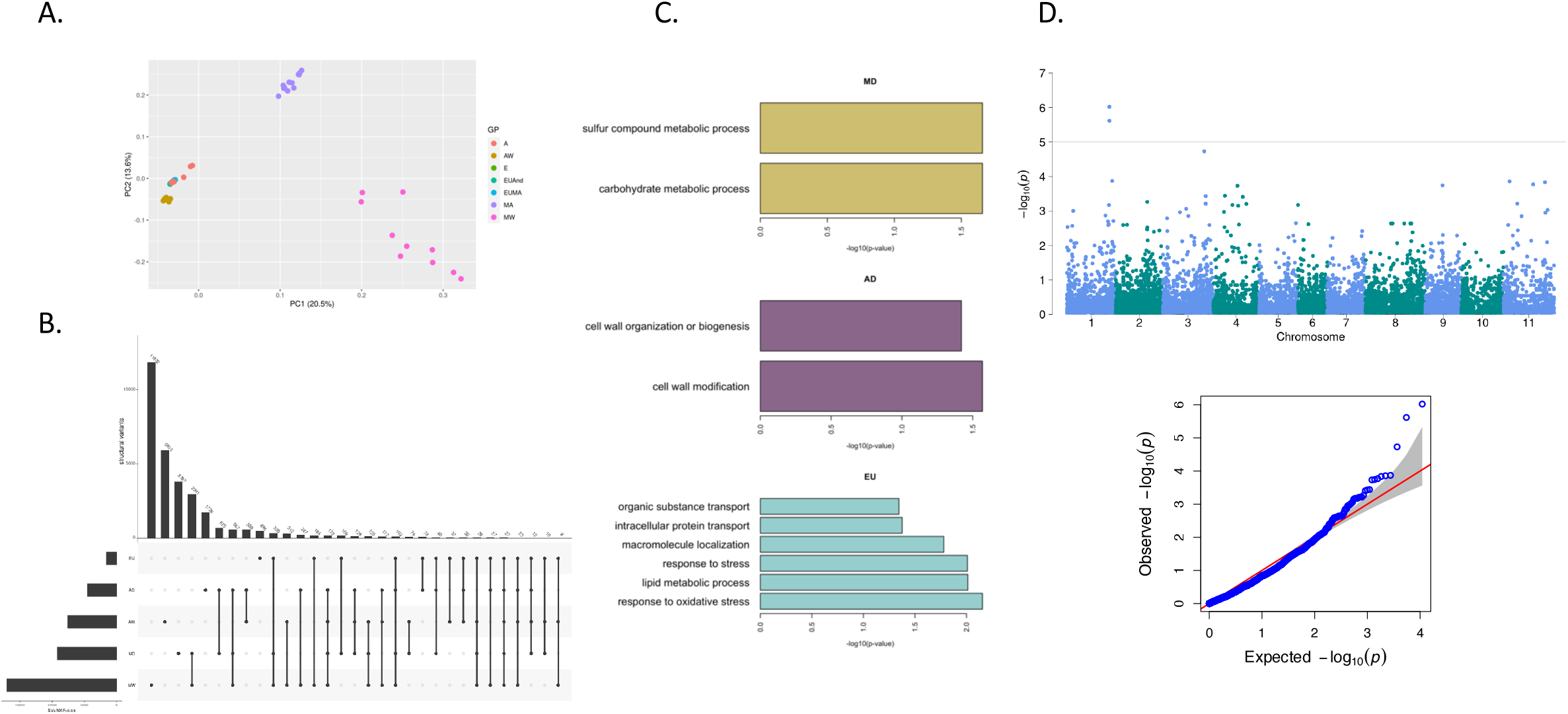
Structural variants. A. PCA based on SVs. B. Overlap of SVs between genepools. C. Functional categories enriched among genes overlapping pool-specific SVs. D. Manhattan plots showing the association of SVs to photoperiod sensitivity.

To further understand the role of SVs in the domestication and adaptation processes we looked for gene models that overlap with gene pool specific predictions of SVs and searched for functional enrichments among those genes. We identified 655, 339 and 255 gene models that overlapped with SVs in the MD, AD and EU pools, respectively, and observed interesting functional categories that were enriched among them (Figure 3C). In the MD pool genes overlapping with SVs were enriched for sulphur compound 368 and carbohydrate metabolism, while the AD and EU gene pools showed enrichments for cell wall modifications and stress response, respectively. Finally, a GWAS based on the SVs identified two variants located on chromosome 1 that were associated with the capacity to flower during Scandinavian summers: an insertion of 66bp at 44,731,098 bp and a deletion of 5064bp at 44,783,344 (figure 3D). These variants are located in the vicinity of *TFL1*.

### Flowering associated haplotypes

We ran genome wide association analyses using both multi-locus models, MLMM and Blink, as well as single locus model, MLM, all implemented in GAPIT3; the advantages of these models in terms of statistical power vs computational cost have been discussed elsewhere (Wang and Zhang 2021). All methods detected a significant signal (Bonferroni correction alpha=0.05, *p*-value<3-1e-7) around 44.8Mb on chromosome 1 (Figure 4A) that matches the previously identified *Fin* locus (Kwak, et al. 2008; Kwak, et al. 2012). At this locus, the highest scoring SNP (SNP Chr01_44852374_G_A, MLMM p-value of 2.8e^-8^) explains 43% of the phenotypic variance and is located in close proximity (less than 4Kb) to *Terminal Flower 1* (*TFL1*, Phvul.001G189200 located between 44,856,139 and 44,857,862bp), which has previously shown to be responsible for the indeterminate phenotype in *Phaseolus vulgaris* (Koinange, et al. 1996). Using extended haplotype homozygosity (EHH) and haplotype length, we observed in flowering accessions an extended EHH (>0.4 after 44.9Mb), whereas accessions that could not flower in long days had a very rapid decay of EHH and most of them carried the ancestral allele.

**Figure 4.**
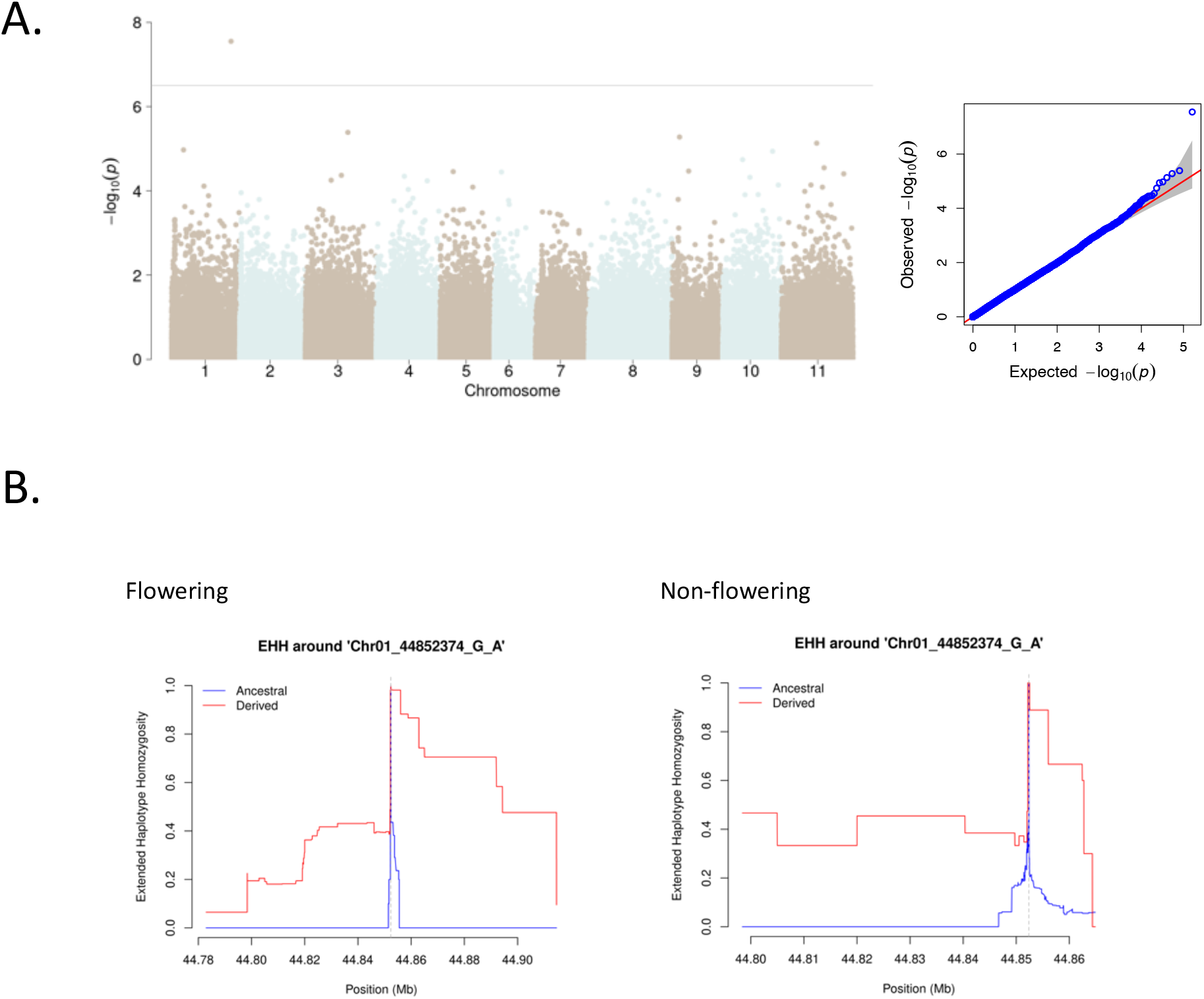
Dissection of photoperiod response. A. GWAS to photoperiod sensitivity in our collection of *P. vulgaris* (MLMM, significance threshold -log10(*p*-value)=6.5). B. Extended homozygosity around TFL1.

## Discussion

The capacity to flower under long days is a fundamental trait for bean cultivation in Scandinavia. During the spring and summer seasons, daylength rapidly transitions form approximately 10 to 17hrs between the months of March and July. Due to the risk of frost damage during early developmental stages, common bean cultivation launches late in the spring and thus, the transition from vegetative to reproductive stages occurs at non-neutral daylengths. For this reason, we evaluated the flower capacity on a large collection of common bean accessions during the summer months in Sweden (at a latitude of 59.8°N). We observed strong GWAS signals (SNPs and SVs) and differences in the extended haplotype homozygosity around a key regulator of photoperiod sensitivity, *TERMINAL FLOWER 1* (*TFL1*), on chromosome 1. *TFL1* is a floral repressor which is closely related to the florigen gene *FLOWERING LOCUS T* (*FT*, (Corbesier, et al. 2007)). Both *TFL1* and *FT* are mobile proteins, but while *FT* moves from the leaves to the shoot apical meristem (SAM), *TFL1* moves within the SAM, regulating flowering time and shoot indeterminacy. *TFL1* is also involved in maintaining the SAM, allowing indeterminate growth of the inflorescence. During the vegetative phase, *TFL1* is expressed in low levels in the centre of SAM, but during the switch to reproductive phase, *TFL1* is strongly up-regulated in axilliary meristems and then in the SAM which then converts into an inflorescence meristem. This locus has been repeatedly associated to indeterminacy in *P. vulgaris* in both the Mesoamerican and Andean gene pools (Kwak, et al. 2008; Kwak, et al. 2012). Furthermore, using QTL mapping on a RIL population derived from Andean x Mesoamercian crosses with determinate and indeterminate genotypes, Gonzalez and collaborators (Gonzalez, et al. 2016) suggested that TFL1 might be also involved in the flowering response and not only defining the indeterminacy of the plants, as the identified QTL around TFL-1 explained up to 32% of phenotypic variation for time to flowering, 66% for vegetative growth, and 19% for rate of plant production.

### Functional convergence during domestication and adaptation

We observe a remarkable parallelism at the functional level in terms of the gene ontology categories that were significantly enriched in both domestication events and during the adaptation of the common bean to European conditions. Given the drastic changes in terms of seed traits, as varieties have been selected for increasing their nutritional value, it was not surprising to find carbohydrate metabolism related categories enriched for both events of artificial selection. However, more unexpected GO terms, such as DNA repair related categories and protein phosphorylation, were also enriched in positive selection outliers in both Andean and Mesoamerican gene pools, suggesting parallel targets of artificial selection during both domestication events. The role of such categories behind domestication has only recently been addressed by Wang and collaborators (Wang, et al. 2019), who suggest that if the accumulation of polymorphisms in DNA repair related genes predates domestication, they could have facilitated the emergence of domestication syndrome traits. In addition, enrichment of protein (de)phosphorylation, the other functional category present for both domestication events, could easily occur due to the important role of post-translational modifications in plant development and, most importantly, in response to different sources of stress (reviewed by (Damaris and Yang 2021)).

### Structural variants behind domestication and adaptation

Improving methods of SV detection has led to an increasing amount of evidence supporting their effect on a wide variety of plant traits. SVs have, for example, been shown to play a role in plant resistance and immunity such as copy number variation (CNV) in the aluminium-resistance gene *MATE1* that affects aluminium tolerance in maize (Maron, et al. 2013), an increase in VRN-A1 locus copy number in wheat that has been associated with frost tolerance (Zhu, et al. 2014) and presence/absence variation (PAV) that has been linked to resistance against *Verticillium longisporum* fungal infection in rapeseed, *Brassica napus* (Gabur, et al. 2019). SVs have also been shown to affect phenotypic plant traits. Fruit shape and flesh colouring have been shown to be associated with a 487 bp DEL and a 1.67 Mb INV respectively in peach (Guo, et al. 2020). Yang and collaborators (Yang, et al. 2019) found DELs and INVs that were significantly associated to oil concentration and long-chain fatty acid composition in maize. Ecotypic differentiation resulting from SV formation has been observed in wild sunflower where divergent haploblocks have further been associated with seed size and flowering time (Todesco, et al. 2020). Also, it was demonstrated that changes in gene dosage and expression levels resulting from SVs in tomato can affect traits such as fruit flavour, size and production (Alonge, et al. 2020).

Here we are able for the first time to associate structural variants to traits that have been important during the domestication process by identifying pool-specific SVs in the common bean. We produced a high-quality catalogue of structural variants that were supported by three independent tools, and this allowed us to evaluate both the proximity of SVs to gene models as well as their implication for the phenotypic traits we have dissected in this report. The functional enrichments of gene models identified as overlapping with SVs and the strong association of two particular SVs to photoperiod sensitivity are compelling arguments that support the relevance of SVs in phenotypic evolution in the common bean. While analyses of gene expression differences could provide even further details about the regulatory role of SVs in the common bean, the associations reported herein provide important cues about the multi-layered genomic landscape of *P. vulgaris*, from its domestication to its recent introduction and adaptation to Europe.

## Acknowledgements

The common bean collection was phenotyped at the Plant Cultivation Facility in Biocentrum, SLU campus Ultuna. We want to thank the team in charge of the phytotrons and green-house, Urban Pettersson, Per Linden, Fredric Hedlund and Kathrin Hesse. This research was financed by the Swedish Research Council (VR), under the grant no. 2018-03780 to PKI. We acknowledge support from the National Genomics Infrastructure in Stockholm funded by Science for Life Laboratory, the Knut and Alice Wallenberg Foundation and the Swedish Research Council for assistance with massively parallel sequencing. All computational analyses and data handling were enabled by resources provided by the Swedish National Infrastructure for Computing (SNIC) at Uppsala Multidisciplinary Centre for Advanced Computational Science (UPPMAX) under the computing projects 2019/3-597, 2020/5-621 and storage project sllstore2017050.

## Data accessibility

Raw read data have been uploaded to NCBI and can be found under the SRA Bioproject PRJNA1004188.

## Author contributions

MR-A and PKI planned and designed the research. MR-A, GB and LY performed experiments and analyzed data. MR-A and PKI wrote the manuscript. All authors read and approved of the final version of the manuscript.

