## Supplementary material for "Haplotype differences in common bean accessions confer the capacity to flower under Scandinavian summer conditions": supp_material

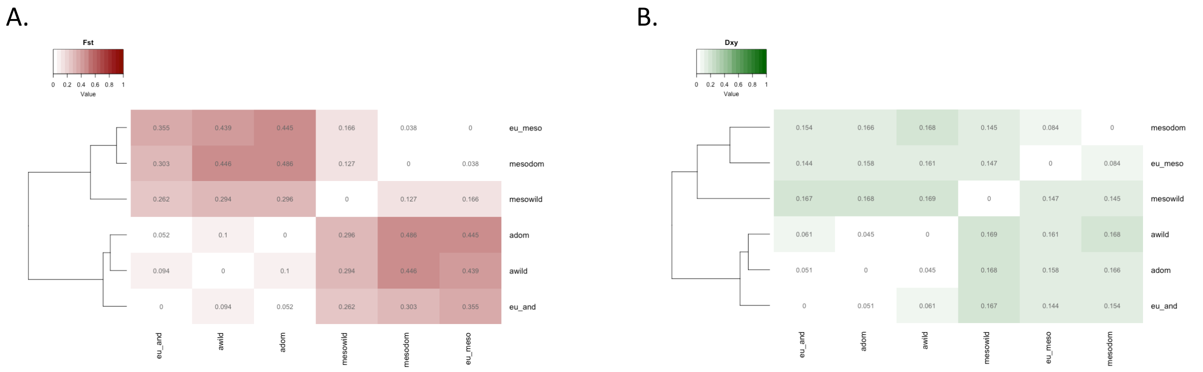


**Supplementary Figure 1.** Differentiation estimators between *P.vulgaris* subpopulations. A. Heatmap of pairwise $F_{ST}$ values. B. Heatmap of pairwise $D_{XY}$ values.

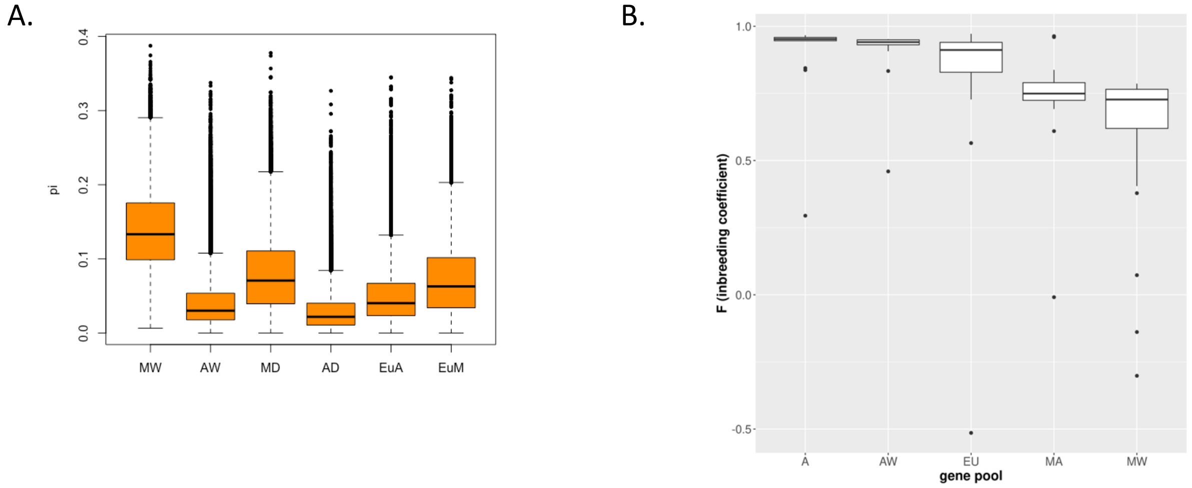


**Supplementary Figure 2.** Diversity estimators. A. Nucleotide diversity per subpopulation. B. Inbreeding coefficient per subpopulation.

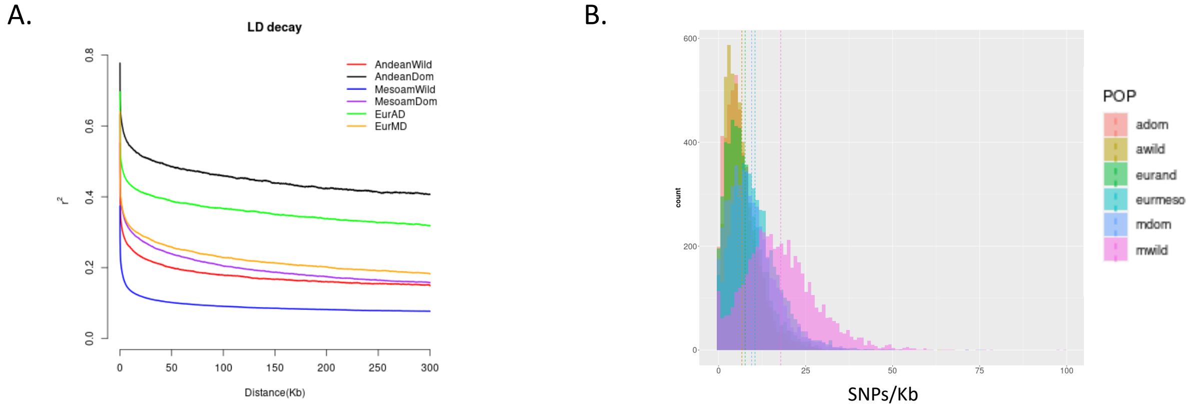


**Supplementary Figure 3.** A. Linkage disequilibrium decay in each subpopulation. B. SNP density per Kb.

**Supplementary Table 1.** *P. vulgaris* accessions used in this study. The seed banks where the material was obtained are the International Center for Tropical Agriculture, Colombia (CIAT; accession ID: Gxxx), the Nordic Genetic Resource Center (NordGen; ID NGBxxx) and European Search Catalogue for Plant Genetic Resources at Gatersleben, Germany (IPK; ID: PHAxxx).

| Seq-ID | Accession | Gene-Pool | Country of Origin | Coverage | |
|---|---|---|---|---|---|
| | | | | **Depth** | **Breadth** |
| **P13861_101** | PHA865 | EU | Italy | 13,18820852 | 0,87569681 |
| **P13861_102** | G2868 | MA | Mexico | 7,34614598 | 0,851759932 |
| **P13861_103** | NGB23858 | EU | Norway | 6,846813622 | 0,898106158 |
| **P13861_104** | PHA49 | EU | Sweden | 7,431964691 | 0,907081044 |
| **P13861_105** | G4338 | MA | Mexico | 5,42535001 | 0,836760801 |
| **P13861_106** | G1461 | EU | Sweden | 6,482791319 | 0,904517851 |
| **P13861_107** | G5340 | EU | Sweden | 7,241256986 | 0,884736296 |
| **P13861_108** | PHA1697 | EU | Poland | 8,041974637 | 0,923147199 |
| **P13861_109** | G18939C | EU | Sweden | 7,656379534 | 0,915632754 |
| **P13861_110** | NGB9300 | EU | Norway | 8,01626446 | 0,905477118 |
| **P13861_111** | G23422 | AW | Peru | 7,341778465 | 0,907401857 |
| **P13861_112** | NGB17811 | EU | Sweden | 7,287214586 | 0,907742447 |
| **P13861_113** | PHA1780 | EU | Spain | 7,983646578 | 0,905108947 |
| **P13861_114** | PHA13545 | EU | Belgium | 8,051885812 | 0,908524456 |
| **P13861_115** | NGB17816 | EU | Sweden | 23,2366404 | 0,917752057 |
| **P13861_116** | PHA1137 | EU | Hungary | 11,87773447 | 0,917438767 |
| **P13861_117** | PHA13960 | EU | Spain | 8,830264889 | 0,909430212 |
| **P13861_118** | G20325 | EU | Sweden | 8,791892891 | 0,909326748 |
| **P13861_119** | NGB18054 | EU | Sweden | 9,904820787 | 0,915633673 |
| **P13861_120** | G11285 | EU | Sweden | 11,42013536 | 0,925896048 |
| **P13861_121** | PHA5866 | EU | Italy | 9,748211974 | 0,907625293 |
| **P13861_122** | G24404 | MW | Colombia | 8,570800122 | 0,84967909 |
| **P13861_123** | NGB4150 | EU | Sweden | 8,183897474 | 0,908515504 |
| **P13861_124** | NGB13466 | EU | Norway | 6,120912829 | 0,844600171 |
| **P13861_125** | PHA366 | EU | Italy | 6,960273124 | 0,919419672 |
| **P13861_126** | G12138 | A | Peru | 6,719789617 | 0,916403369 |
| **P13861_127** | NGB17813 | EU | Sweden | 6,694101297 | 0,919555838 |
| **P13861_128** | G23455 | AW | Peru | 6,952608513 | 0,91584377 |
| **P13861_129** | PHA13184 | EU | Slovakia | 7,082909981 | 0,91659738 |
| **P13861_130** | G23423 | AW | Peru | 7,074014253 | 0,902487054 |
| **P13861_131** | G23589 | AW | Peru | 6,470189067 | 0,902085351 |
| **P13861_132** | G13614 | MA | Mexico | 14,66117901 | 0,87970335 |
| **P13861_133** | PHA307 | EU | Albania | 11,25448217 | 0,867375668 |

| | | | | | |
|---|---|---|---|---|---|
| **P13861_134** | NGB20124 | EU | Denmark | 19,53307721 | 0,922821387 |
| **P13861_135** | G12947 | MW | Mexico | 10,27319731 | 0,876780473 |
| **P13861_136** | PHA6155 | EU | United Kingdom | 13,55195658 | 0,90851483 |
| **P13861_137** | G23418 | MW | Costa Rica | 10,87681216 | 0,858057854 |
| **P13861_138** | G13948 | A | Argentina | 31,34678265 | 0,931955972 |
| **P13861_139** | G19893 | AW | Argentina | 14,32948711 | 0,921733625 |
| **P13861_140** | G12856 | AW | Peru | 14,67828737 | 0,903530603 |
| **P13861_141** | G8982 | EU | Sweden | 18,50969458 | 0,922872542 |
| **P13861_142** | PHA13186 | EU | Slovakia | 18,18522843 | 0,932968104 |
| **P13861_143** | G9836 | A | Bolivia | 16,82092818 | 0,932052098 |
| **P13861_144** | PHA13736 | EU | United Kingdom | 17,53583884 | 0,904211422 |
| **P13861_145** | NGB17807 | EU | Sweden | 17,80676871 | 0,916122644 |
| **P13861_146** | PHA1887 | EU | Poland | 12,49317197 | 0,905225639 |
| **P13861_147** | G23578A | MA | Colombia | 11,16156867 | 0,871730079 |
| **P13861_148** | PHA13609 | EU | Switzerland | 14,90622099 | 0,929328209 |
| **P13861_149** | NGB21237 | EU | Sweden | 18,7546417 | 0,924120114 |
| **P13861_150** | PHA2680 | EU | Sweden | 20,54461505 | 0,922054169 |
| **P13861_151** | G1460 | EU | Sweden | 17,83240401 | 0,890759428 |
| **P13861_152** | NGB11569 | EU | Norway | 20,15032933 | 0,925208996 |
| **P13861_153** | PHA13666 | EU | United Kingdom | 19,11561485 | 0,895566851 |
| **P13861_154** | G4534 | A | Colombia | 12,38351074 | 0,925607153 |
| **P13861_155** | NGB23934 | EU | Sweden | 11,911512 | 0,913970269 |
| **P13861_156** | NGB1828 | EU | Denmark | 9,22269806 | 0,909258677 |
| **P13861_157** | G13094 | MA | Mexico | 13,73027023 | 0,928476014 |
| **P13861_158** | G10075 | EU | Netherlands | 13,91733087 | 0,925682811 |
| **P13861_159** | G1281 | EU | Sweden | 11,9558986 | 0,864293999 |
| **P13861_160** | G24408 | MW | Colombia | 12,16215319 | 0,870573544 |
| **P13861_161** | G8697 | A? | Bolivia | 15,03347171 | 0,928096984 |
| **P13861_162** | G18939B | EU | Sweden | 14,11838142 | 0,896958616 |
| **P13861_163** | G9456 | EU | Sweden | 10,64150331 | 0,913012921 |
| **P13861_164** | PHA13061 | EU | Belgium | 10,63760361 | 0,910218654 |
| **P13861_165** | G18939A | EU | Sweden | 12,04109998 | 0,893931554 |
| **P13861_166** | G8811 | EU | Sweden | 20,05433272 | 0,898412709 |
| **P13861_167** | PHA4534 | EU | Hungary | 11,49516786 | 0,909150471 |
| **P13861_168** | PHA1772 | EU | Slovakia | 9,640424652 | 0,860314797 |
| **P13861_169** | PHA332 | EU | Austria | 9,266253621 | 0,862349126 |
| **P13861_170** | G11287 | EU | Sweden | 15,23164469 | 0,916021013 |
| **P13861_171** | G8658 | EU | Sweden | 13,08866556 | 0,911499198 |
| **P13861_172** | NGB11573 | EU | Sweden | 13,32917863 | 0,9281691 |
| **P13861_173** | G1540 | EU | Sweden | 12,66576809 | 0,915587683 |
| **P13861_174** | G8763 | EU | Sweden | 17,62148254 | 0,915660055 |
| **P13861_175** | G4383 | MA | Mexico | 11,02610963 | 0,870085711 |

| | | | | | |
|---|---|---|---|---|---|
| **P13861_176** | G23459 | AW | Peru | 11,58838802 | 0,911928364 |
| **P13861_177** | PHA1072 | EU | Switzerland | 10,62386113 | 0,909902245 |
| **P13861_178** | G23419A | AW | Peru | 21,92693302 | 0,919346413 |
| **P13861_179** | G900 | EU | Sweden | 17,23603429 | 0,874269297 |
| **P13861_180** | PHA6437 | EU | Spain | 21,76759146 | 0,921540866 |
| **P13861_181** | PHA1022 | EU | Poland | 18,29886907 | 0,920940453 |
| **P13861_182** | PHA1138 | EU | Hungary | 21,43508304 | 0,919433569 |
| **P13861_183** | PHA725 | EU | Italy | 22,0898583 | 0,924406111 |
| **P13861_184** | PHA368 | EU | Italy | 19,76312672 | 0,877409902 |
| **P13861_185** | PHA2899 | EU | Austria | 20,18491336 | 0,902376497 |
| **P13861_186** | G23426 | AW | Peru | 15,05809279 | 0,912440229 |
| **P13861_187** | NGB23936.3 | EU | Sweden | 14,62018133 | 0,92108872 |
| **P13861_188** | PHA1076 | EU | Hungary | 15,14486729 | 0,921246457 |
| **P13861_189** | NGB17817.5 | EU | Sweden | 21,96682077 | 0,923913125 |
| **P13861_190** | NGB11547.2 | EU | Sweden | 12,95289415 | 0,927020581 |
| **P13861_191** | NGB13468.2 | EU | Sweden | 15,49076553 | 0,929675353 |
| **P13861_192** | PHA3620 | EU | Austria | 24,37764174 | 0,922391899 |
| **P13861_193** | G22303 | MW | Colombia | 17,61702732 | 0,871413573 |
| **P13861_194** | PHA2682 | EU | Sweden | 21,33541001 | 0,92000416 |
| **P13861_195** | NGB17826.5 | EU | Sweden | 21,80600495 | 0,928186147 |
| **P13861_196** | NGB23859.3 | EU | Norway | 18,98924615 | 0,917438389 |
| **P13861_201** | NGB20200.4 | EU | Sweden | 11,75687092 | 0,929154448 |
| **P13861_202** | PHA13112 | EU | United Kingdom | 13,65022882 | 0,914158473 |
| **P13861_203** | G23604A | A | Peru | 10,61455957 | 0,927214656 |
| **P13861_204** | G10093 | EU | Netherlands | 8,789730258 | 0,92115331 |
| **P13861_205** | G1282 | EU | Sweden | 12,78462824 | 0,928022026 |
| **P13861_206** | PHA14276 | EU | Switzerland | 9,926910717 | 0,912982723 |
| **P13861_207** | PHA13035 | EU | Italy | 13,98312537 | 0,925785601 |
| **P13861_208** | PHA361 | EU | Poland | 10,12572385 | 0,922895856 |
| **P13861_209** | G4681 | MA | Colombia | 12,78432014 | 0,935644392 |
| **P13861_210** | G12949 | MW | Mexico | 12,13717757 | 0,87617708 |
| **P13861_211** | G23447 | A | Bolivia | 13,94715036 | 0,946792012 |
| **P13861_212** | PHA13187 | EU | Slovakia | 11,82958707 | 0,870955787 |
| **P13861_213** | PHA722 | EU | Italy | 14,77480974 | 0,884597883 |
| **P13861_214** | PHA13928 | EU | Switzerland | 14,68414825 | 0,911621688 |
| **P13861_215** | G21201 | AW | Argentina | 13,6225733 | 0,917668007 |
| **P13861_216** | PHA6389 | EU | Romania | 11,09127563 | 0,91443794 |
| **P13861_217** | G8920 | MA | Mexico | 14,15216613 | 0,873838572 |
| **P13861_218** | G11035 | MA | Mexico | 11,20903374 | 0,866273956 |
| **P13861_219** | PHA6254 | EU | Albania | 13,83000737 | 0,922984824 |
| **P13861_220** | PHA7313 | EU | Poland | 11,16514748 | 0,920893545 |
| **P13861_221** | PHA13181 | EU | Slovakia | 15,77309223 | 0,900907514 |

| | | | | | |
|---|---|---|---|---|---|
| **P13861_222** | PHA4008 | EU | Italy | 10,02563007 | 0,866937863 |
| **P13861_223** | PHA339 | EU | Italy | 12,27813634 | 0,871921374 |
| **P13861_224** | NGB24077.2 | EU | Sweden | 12,21480718 | 0,871979955 |
| **P13861_225** | PHA12283 | EU | United Kingdom | 11,71705806 | 0,914750874 |
| **P13861_226** | NGB24038.3 | EU | Sweden | 13,93427704 | 0,926027628 |
| **P13861_227** | PHA167 | EU | Greece | 12,30593846 | 0,918596675 |
| **P13861_228** | PHA109 | EU | Greece | 9,840950799 | 0,916548558 |
| **P13861_229** | G16310 | MA | Guatemala | 11,58422351 | 0,868671596 |
| **P13861_230** | NGB11751.2 | EU | Sweden | 12,25938798 | 0,917263535 |
| **P13861_231** | NGB17814.3 | EU | Sweden | 16,39002343 | 0,915168092 |
| **P13861_232** | NGB23840.2 | EU | Sweden | 12,1808214 | 0,871372575 |
| **P13861_233** | PHA1753 | EU | Romania | 12,27223604 | 0,917206832 |
| **P13861_234** | G19889 | AW | Argentina | 11,00733436 | 0,918894543 |
| **P13861_235** | PHA1086 | EU | Belarus | 10,62967617 | 0,916692961 |
| **P13861_236** | PHA12934 | EU | Italy | 12,7905154 | 0,874300952 |
| **P13861_237** | PHA5934 | EU | Albania | 19,83813127 | 0,93080588 |
| **P13861_238** | PHA143 | EU | Poland | 15,26267389 | 0,930148953 |
| **P13861_239** | PHA1077 | EU | Belgium | 13,73052998 | 0,915413558 |
| **P13861_240** | G23445 | AW | Bolivia | 10,15263243 | 0,914981432 |
| **P13861_241** | G6762 | MA | Mexico | 11,75124912 | 0,872438571 |
| **P13861_242** | G23434A | MA | Guatemala | 10,04118271 | 0,88798853 |
| **P13861_243** | G23444 | AW | Bolivia | 16,2762085 | 0,916484904 |
| **P13861_244** | G22033 | MA | Mexico | 11,66443543 | 0,86874677 |
| **P13861_245** | G7229 | A? | Colombia | 10,64265362 | 0,929366011 |
| **P13861_246** | PHA13099 | EU | United Kingdom | 14,2779352 | 0,910339475 |
| **P13861_247** | PHA3669 | EU | Austria | 11,38965227 | 0,92563513 |
| **P13861_248** | PHA13612 | EU | Sweden | 10,3563034 | 0,913315682 |
| **P13861_249** | G7648 | EU | Sweden | 16,41090415 | 0,917838801 |
| **P13861_250** | PHA1142 | EU | Hungary | 19,55906961 | 0,917174552 |
| **P13861_251** | G21069 | A | Argentina | 15,19180914 | 0,927479372 |
| **P13861_252** | G21197 | AW | Argentina | 22,16943967 | 0,922497095 |
| **P13861_253** | PHA419 | EU | Switzerland | 19,98231616 | 0,913728252 |
| **P13861_254** | PHA7138 | EU | Spain | 15,99687402 | 0,876078979 |
| **P13861_255** | PHA841 | EU | Switzerland | 15,93969258 | 0,911399868 |
| **P13861_256** | PHA7309 | EU | Poland | 19,51058229 | 0,925714412 |
| **P13861_257** | G3296 | MA | Mexico | 11,26489342 | 0,8750718 |
| **P13861_258** | G23777 | A | Peru | 12,88936473 | 0,932524891 |
| **P13861_259** | NGB17824 | EU | Sweden | 12,00503806 | 0,915874923 |
| **P13861_260** | PHA3673 | EU | Austria | 15,34498649 | 0,931658061 |
| **P13861_261** | PHA6066 | EU | Italy | 11,74428632 | 0,923422939 |
| **P13861_262** | G21194 | AW | Argentina | 15,93513057 | 0,923274788 |
| **P13861_263** | PHA1752 | EU | Romania | 9,832059789 | 0,866373591 |

| | | | | | |
|---|---|---|---|---|---|
| **P13861_264** | PHA3687 | EU | Austria | 11,4497756 | 0,913309476 |
| **P13861_265** | G23458 | AW | Peru | 12,08971139 | 0,916122351 |
| **P13861_266** | G18939 | EU | Sweden | 11,91040824 | 0,906361549 |
| **P13861_267** | PHA306 | EU | Albania | 8,264037238 | 0,853386069 |
| **P13861_268** | G10000 | MA | Mexico | 10,25386419 | 0,868538416 |
| **P13861_269** | NGB24316.1 | EU | Sweden | 9,175246151 | 0,86711039 |
| **P13861_270** | PHA7150 | EU | Spain | 9,629021496 | 0,9062057 |
| **P13861_271** | G15914 | EU | Netherlands | 8,686600999 | 0,866603379 |
| **P13861_272** | G23464 | MW | Mexico | 8,998508445 | 0,852600432 |
| **P13861_273** | PHA6287 | EU | Albania | 20,75423559 | 0,883733211 |
| **P13861_274** | NGB18415.1 | EU | Sweden | 14,02765672 | 0,907186213 |
| **P13861_275** | PHA1450 | EU | Slovakia | 14,94152033 | 0,879897761 |
| **P13861_276** | PHA154 | EU | Greece | 12,96804481 | 0,874731431 |
| **P13861_277** | G10966 | MA | Mexico | 16,48832482 | 0,876912124 |
| **P13861_278** | PHA158 | EU | Poland | 19,13090236 | 0,889941755 |
| **P13861_279** | NGB23857.4 | EU | Denmark | 20,73102752 | 0,918857664 |
| **P13861_280** | NGB28356.3 | EU | Sweden | 20,53657437 | 0,926821448 |
| **P13861_281** | PHA5989 | EU | Romania | 23,79099251 | 0,924107458 |
| **P13861_282** | G16843 | A | Peru | 19,9130139 | 0,933487471 |
| **P13861_283** | NGB1829 | EU | Denmark | 16,50172423 | 0,922395529 |
| **P13861_284** | PHA4620 | EU | Hungary | 24,52186404 | 0,915719147 |
| **P13861_285** | G13177 | MA | Mexico | 13,78359717 | 0,878813625 |
| **P13861_286** | G11059 | MA | Mexico | 15,78399704 | 0,88206065 |
| **P13861_287** | G24345 | MW | Mexico | 14,31962146 | 0,87614694 |
| **P13861_288** | G14629 | EU | Sweden | 13,89553687 | 0,9017849 |
| **P13861_289** | PHA14277 | EU | Austria | 11,21685484 | 0,863497379 |
| **P13861_290** | G20324 | EU | Sweden | 19,00915468 | 0,877540836 |
| **P13861_291** | G57 | A | Canada | 21,61786058 | 0,931238735 |
| **P13861_292** | G21056 | A | Argentina | 29,71249843 | 0,933341886 |
| **P13861_293** | G11053 | MW | Mexico | 9,582526956 | 0,882638165 |
| **P13861_294** | G1683 | EU | Sweden | 21,65900497 | 0,904290965 |
| **P13861_295** | G21043 | A | Argentina | 14,483144 | 0,931162909 |
| **P13861_296** | G11015 | MA | Mexico | 16,67883854 | 0,874468085 |
| **P13861_301** | PHA1139 | EU | Hungary | 7,250160054 | 0,869941489 |
| **P13861_302** | G23435 | MW | Guatemala | 10,40022539 | 0,859942372 |
| **P13861_303** | PHA244 | EU | Poland | 11,08118475 | 0,901009938 |
| **P13861_304** | PHA13188 | EU | Slovakia | 9,243250417 | 0,901645689 |
| **P13861_305** | G23556 | MW | Mexico | 9,813465092 | 0,866071467 |
| **P13861_306** | G24322 | AW | Argentina | 10,87322183 | 0,916402304 |
| **P13861_307** | G24318 | AW | Argentina | 12,42394236 | 0,915708211 |
| **P13861_308** | G5341 | EU | Sweden | 10,32815272 | 0,863881677 |
| **P13861_309** | G24605 | MW | Mexico | 10,27426934 | 0,863694308 |

| | | | | | |
|---|---|---|---|---|---|
| **P13861_310** | PHA13183 | EU | Slovakia | 10,00010527 | 0,864887357 |
| **P13861_311** | G24412 | MW | Colombia | 9,31672015 | 0,93468336 |
| **P13861_312** | PHA13224 | EU | Albania | 8,912686087 | 0,855113913 |
| **P13861_313** | NGB23839.2 | EU | Sweden | 11,95551927 | 0,928925498 |
| **P13861_314** | PHA920 | EU | Romania | 8,197108129 | 0,856394898 |
| **P13861_315** | G23421 | AW | Peru | 8,373524354 | 0,9041143 |
| **P13861_316** | PHA6011 | EU | Romania | 10,74237596 | 0,909507123 |
| **P13861_317** | G12875 | MW | Mexico | 10,98649997 | 0,868804252 |
| **P13861_318** | G19898 | AW | Argentina | 8,271373641 | 0,912003475 |
| **P13861_319** | PHA987 | EU | Hungary | 7,492220199 | 0,850654842 |
| **P13861_320** | PHA99 | EU | Greece | 5,855819432 | 0,837518831 |
| **P13861_321** | G18939D | EU | Sweden | 9,868897569 | 0,903235388 |
| **P13861_322** | NGB23850 | EU | Denmark | 9,025085303 | 0,908526915 |
| **P13861_323** | G23442 | AW | Bolivia | 8,518639735 | 0,904744982 |
| **P13861_324** | PHA5877 | EU | Italy | 21,11910522 | 0,919093559 |
| **P13861_325** | PHA13189 | EU | Slovakia | 12,81074329 | 0,86934349 |
| **P13861_326** | PHA14278 | EU | Austria | 17,90032327 | 0,925025306 |
| **P13861_327** | G24390 | MW | Mexico | 21,04494639 | 0,873518301 |
| **P13861_328** | PHA295 | EU | Albania | 23,11331545 | 0,884057684 |
| **P13861_330** | PHA5909 | EU | Albania | 19,4598115 | 0,8783244 |
| **P13861_331** | G22837 | MW | Mexico | 26,57550759 | 0,886558189 |
| **P13861_332** | G12873 | MW | Mexico | 20,18867228 | 0,884560738 |
| **P13861_333** | G13955 | A | Argentina | 19,9833669 | 0,929909965 |
| **P13861_334** | G7930 | A | Argentina | 21,20021355 | 0,900950731 |
| **P13861_335** | G23457A | A | Peru | 17,81185037 | 0,929376372 |
| **P13861_336** | G12879 | MW | Mexico | 18,74383879 | 0,891219788 |
| **P13861_337** | G24323 | MW | Mexico | 16,72270408 | 0,870774276 |
| **P13861_338** | G12865 | MW | Mexico | 17,48735604 | 0,87011042 |
| **P13861_339** | PHA7686 | EU | Romania | 15,17963105 | 0,869685937 |
| **P13861_340** | PHA13228 | EU | Slovakia | 20,91318455 | 0,928909637 |

**Supplementary table 2.** Classification of European cultivars according to their genomic background.

| EU-Andean background | | EU-Mesoamerican background |
|---|---|---|
| EU_G10075 | EU_PHA14278 | EU_G1281 |
| EU_G10093 | EU_PHA143 | EU_G1460 |
| EU_G1282 | EU_PHA167 | EU_G14629 |
| EU_G1683 | EU_PHA1697 | EU_G15914 |
| EU_G18939 | EU_PHA1753 | EU_G5341 |
| EU_G18939D | EU_PHA1780 | EU_G900 |
| EU_G5340 | EU_PHA1887 | EU_NGB24316 |
| EU_G8658 | EU_PHA244 | EU_PHA1139 |
| EU_NGB13468 | EU_PHA2682 | EU_PHA12934 |
| EU_NGB17826 | EU_PHA2899 | EU_PHA13666 |
| EU_NGB18415 | EU_PHA361 | EU_PHA14277 |
| EU_NGB20124 | EU_PHA3620 | EU_PHA154 |
| EU_NGB23857 | EU_PHA366 | EU_PHA158 |
| EU_NGB23858 | EU_PHA3669 | EU_PHA1772 |
| EU_NGB23934 | EU_PHA3673 | EU_PHA295 |
| EU_NGB23936 | EU_PHA3687 | EU_PHA307 |
| EU_NGB24038 | EU_PHA419 | EU_PHA332 |
| EU_NGB9300 | EU_PHA4534 | EU_PHA339 |
| EU_PHA1022 | EU_PHA4620 | EU_PHA4008 |
| EU_PHA1076 | EU_PHA49 | EU_PHA5909 |
| EU_PHA1077 | EU_PHA5866 | EU_PHA6287 |
| EU_PHA1086 | EU_PHA5877 | EU_PHA722 |
| EU_PHA109 | EU_PHA5934 | EU_PHA7686 |
| EU_PHA1137 | EU_PHA5989 | EU_PHA865 |
| EU_PHA1138 | EU_PHA6011 | EU_PHA987 |
| EU_PHA1142 | EU_PHA6066 | EU_PHA99 |
| EU_PHA13035 | EU_PHA6155 | |
| EU_PHA13099 | EU_PHA6254 | |
| EU_PHA13112 | EU_PHA6389 | |
| EU_PHA13184 | EU_PHA6437 | |
| EU_PHA13188 | EU_PHA7150 | |
| EU_PHA13228 | EU_PHA725 | |
| EU_PHA13609 | EU_PHA7309 | |
| EU_PHA13736 | EU_PHA7313 | |
| EU_PHA13928 | EU_PHA841 | |
| EU_PHA13960 | | |
